# Can we predict sleep health based on brain features? A large-scale machine learning study using UK Biobank

**DOI:** 10.1101/2024.10.13.618080

**Authors:** Federico Raimondo, Hanwen Bi, Vera Komeyer, Jan Kasper, Sabrina Primus, Felix Hoffstaedter, Synchon Mandal, Laura Waite, Juliane Winkelmann, Konrad Oexle, Simon B. Eickhoff, Masoud Tahmasian, Kaustubh R. Patil

**Affiliations:** Institute of Neuroscience and Medicine, Brain and Behavior (INM-7), Forschungszentrum Jülich, Jülich, Germany; Institute for Systems Neuroscience, Medical Faculty, Heinrich-Heine University Düsseldorf, Düsseldorf, Germany; Institute of Neurogenomics (ING), Helmholtz Zentrum München, Münich, Germany; Department of Nuclear Medicine, University Hospital and Medical Faculty, University of Cologne, Cologne, Germany; Department of Biology, Faculty of Mathematics and Natural Sciences, Heinrich Heine University Düsseldorf, Germany

**Keywords:** Sleep health, grey matter volume, white matter, functional MRI, UK Biobank, machine learning

## Abstract

**Backgrounds:** Several correlational or group comparison evidence highlighted robust associations between sleep health and macro-scale brain organization. However, inter-individual variability is critical in such interplay. Therefore, in this study, we aimed to investigate the role of brain imaging features in predicting diverse sleep health-related characteristics at the individual subject level using the Machine Learning (ML) approach.

**Methods:** A sample of 28,088 participants from the UK Biobank was employed to calculate 4677 structural and functional neuroimaging markers. Then, we employed them to predict self-reported insomnia symptoms, sleep duration, easiness of getting up in the morning, chronotype, daily nap, daytime sleepiness, and snoring. To assess the predictability of brain features, we built seven different linear and nonlinear ML models for each sleep health-related characteristic.

**Results:** We performed extensive ML analyses that involved more than 19 years of compute time. We observed relatively low performance in predicting all sleep health-related characteristics from brain images (e.g., balanced accuracy ranging between 0.50-0.59). Across all models, the best performance achieved was 0.59, using a linear ML model to predict the ease of getting up in the morning. In fact, a similar performance was achieved with models trained solely on age and sex, indicating that these demographic factors might be the ones driving the predictions.

**Conclusions:** The low capability of multimodal neuroimaging markers in predicting sleep health-related characteristics, even under extensive ML optimization in a large population sample, suggests a complex relationship between sleep health and brain organization.

## Introduction

Sleep is a non-negotiable human need, which has pivotal impacts on memory processing, metabolite clearance, immune system adaptation, optimal cognition, and mental health^1^. The intricate relationship between sleep health and brain health has recently garnered significant scientific attention^2–8^. Sleep health (SH) is a multidimensional concept characterized by subjective satisfaction, alertness, regularity, timing, and sleep duration^9^, which is considered a crucial indicator of human well-being. Seven different SH-related characteristics (i.e., sleep duration, easiness/difficulty of getting up in the morning, chronotype, nap, daytime dozing/sleepiness, as well as insomnia symptoms and snoring) reflect various SH dimensions and were collected in half a million participants in the UK Biobank (UKB)^10,11^. This large-scale population data presents a unique opportunity to explore the link between various SH dimensions and brain structure/function, overcoming the low reproducibility of previous small sample studies, as employed previously^3,5,12–15^.

The link between various SH dimensions and brain structure and function has been reported in correlational or group comparison studies but pointed to heterogeneous results. Sleep disturbance conditions, including insomnia symptoms^8,16,17^, sleep-disordered breathing^18–21^, and abnormal sleep duration^5,22^, provided a non-reproducible and inconclusive association between sleep and the brain. For example, *i)* for insomnia domain, Schiel et al. utilized data from the general population in the UKB, while Weihs et al. analyzed both the general population and patients with clinical insomnia disorder from the ENIGMA-Sleep datasets. Neither study found a strong association between insomnia symptoms/disorder and gray matter volume (GMV)^8,23^. However, Stolicyn and colleagues using UKB showed that insomnia symptoms are associated with higher global GMV, mainly in the amygdala, hippocampus, and putamen^24^. Moreover, individuals with insomnia symptoms demonstrated altered functional connectivity (FC) within and between the default mode network (DMN), frontoparietal network (FPN), and salience network (SN)^17^; *ii)* regarding sleep duration, one study using UKB data found that short sleep duration is linked with lower amygdala reactivity to a negative facial expression task^25^. The non-linear associations have been documented between sleep duration, cognitive performance, mental health, and a wide range of regional differences in brain structure, mainly in the subcortical areas^5,23,24,26,27^. Fjell and colleagues performed cross-sectional analyses based on the UKB sample, indicating inverse U-shaped relationships between sleep duration and brain structure, i.e., 6.5 hours of sleep was associated with increased cortical thickness and subcortical volumes relative to intracranial volume. However, they failed to identify a longitudinal association between sleep duration and cortical thickness^4^. In another study, they found that individuals who reported short sleep without other sleep problems or daytime sleepiness had larger brain volumes compared to both short sleepers with sleep issues and daytime sleepiness, as well as those who slept 7−8 hours^28^; *iii)* analysis of chronotypes also showed that the evening chronotype is linked with higher GMV in the precuneus, bilateral nucleus accumbens, caudate, putamen and thalamus, and orbitofrontal cortex^29^. Another study observed an association between chronotype and neuroimaging phenotypes, which is mediated by genetic factors^30^; *iv)* self-reported daytime sleepiness has been reported to be related to higher cortical GMV^31^. These findings together represent an overall inconsistency in the relationship between insomnia, sleep duration, chronotype, and daytime sleepiness with the brain structure and function. While these studies provided valuable insights into the link between each SH dimension and the brain, they were mostly case-control or correlational studies and might not have been able to capture the complex interplay between the brain and SH^32^, which is a heterogeneous subjective concept that varies across individuals. Thus, the substantial inter-individual variability of SH and the differential associations between various SH characteristics and brain measurements necessitate large-scale datasets and more advanced computational approaches to better model this complexity^33,34^, ultimately improving our understanding of the behavioral consequences of sleep-brain interaction.

Machine learning (ML) offers a powerful tool to unravel complex relationships, providing a more nuanced representation than traditional statistical approaches, which is critical in personalized treatment in sleep medicine^35,36^. ML models can consider complex multivariate linear and nonlinear relations to make brain-behavior predictions on unseen brain imaging data and have the potential to identify generalizable patterns in SH-related neurobiology at the individual subject level^37–39^, surpassing conventional group comparisons and correlations. In particular, nonlinear models are necessary to capture sleep duration-brain interplay. Accurate predictive models can contribute to refining our theoretical understanding of the SH-brain relationship. This might pave the way for developing more effective clinical strategies to enable personalized interventions and treatments ^40^. Directional genetic analyses using Mendelian randomization demonstrated that altered SH dimensions are more a consequence than a cause of brain abnormalities^41^. Thus, this study employed large-scale neuroimaging data from the UKB, assessing whether and how multimodal brain measurements (i.e., structural markers including GMV, surface-based morphometric features, as well as intrinsic functional imaging markers of local correlation (LCOR), global correlation (GCOR), and fractional amplitude of low-frequency fluctuations (fALFF)) can differentiate between well-separated conditions in each SH-related characteristic (e.g., differentiating individuals usually having insomnia symptoms from individuals without insomnia symptoms).

## Methods

### Participants

We selected the data of the first imaging visit (instance 2) from the UKB (http://www.ukbiobank.ac.uk), recorded from 2014 onwards at three different sites in the UK (Cheadle, Reading, Newcastle). The acquisition parameters and protocol of both the structural and functional MRI are as described previously^11^. We included all individuals who participated in the imaging session, and their data had already been preprocessed and denoised by the UKB team^42^. Thus, no particular in-/exclusion criteria have been applied in this sample to be representative of the general population. We selected the individuals for which all the features were computed, resulting in a total N of 28,088, 47% male and 64.1 years old on average (58 – 78 years IQR), were included. The UKB project is approved by the NHS National Research Ethics Service (Ref. 11/NW/0382), and all participants gave written informed consent before participation. Ethical standards are continuously controlled by a Ethics Advisory Committee (EAC, http://www.ukbiobank.ac.uk/ethics), based on a project-specific Ethics and Governance Framework (http://www.ukbiobank.ac.uk/wp-content/uploads/2011/05/EGF20082.pdf). The current analyses were conducted under UK Biobank application number 41655.

### Sleep health characteristics

The multifaceted definition of SH in UKB is based on previous SH studies^3,9,17,23,25,43^. Accordingly, the seven SH-related characteristics were self-reported insomnia symptoms, sleep duration, difficulty/easiness of getting up in the morning, chronotype, daily nap, daytime sleepiness, and snoring (category 100057), obtained from the touchscreen questionnaire. As these questions were asked at every visit, we selected the responses from the visit matching the neuroimaging acquisition visit.

- Sleeplessness/insomnia field (field 1200): “Do you have trouble falling asleep at night or do you wake up in the middle of the night?”, which could be answered as “never/rarely”, “sometimes”, “usually” or “prefer not to answer”.
- Sleep duration (field 1160): “How many hours sleep do you get in every 24 hours?”.
- Getting up in the morning (field 1170): “On average a day, how easy do you find getting up in the morning?”, with four answers spanning from not at all easy to very easy, as well as “do not know” and “prefer not to answer”.
- Chronotype (i.e., morning/evening person, field 1180): “What do you consider yourself to be?”, with four possible answers spanning from a “morning person” to an “evening person”, as well as “do not know” and “prefer not to answer”.
- Nap during the day (field 1190): “Do you have a nap during the day?”, which can be answered as “never/rarely”, “sometimes”, “usually” or “prefer not to answer”.
- Daytime dozing (field 1220): “How likely are you to doze off or fall asleep during the daytime when you don’t mean to? (e.g. when working, reading or driving)”, which can be answered as “never/rarely”, “sometimes”, “often” or “prefer not to answer”.
- Snoring (field 1210): “Does your partner or a close relative or friend complain about your snoring?”, with “yes”, “no”, “do not know” and “prefer not to answer” as possible answers.

Given the ambiguous meaning that some questions, and consequently the respective answers, potentially have in the UKB data (e.g. “sometimes” vs “often”), and to simplify the multiclass/continuous target problems into binary classification problems, we first analyzed the performance of models aimed at distinguishing the extreme answers of each SH-related characteristic. In the case of the continuous answer regarding sleep duration in hours, we split the distribution into four quantiles, selecting the first and fourth quantiles as two classes. However, given the concentration of answers around the median (7 hours), this resulted in discarding only the samples that replied 7 hours. The rationale behind considering the extreme values as class labels is to simplify the classification task, resulting in higher predictive performance if there is indeed a relationship between brain imaging data and each SH-related characteristic. A description of the considered answers for each question, as well as the number of samples for each class, can be seen in Table 1.

**Table 1.** List of answers used for each SH-related characteristic to convert the ambiguous answers into binary classification problems. *denotes the questions for which no samples were dropped.

| Sleep health-related characteristic | Extreme Values |  |  |  |
| --- | --- | --- | --- | --- |
|  | Class 0 |  | Class 1 |  |
|  | Answer(s) | #Samples | Answer(s) | #Samples |
| **Insomnia symptoms** | “Never/rarely” | 6127 | “Usually” | 8846 |
| **Sleep duration** | 1<sup>st</sup> quantile [0-6] | 6760 | 4<sup>th</sup> quantile [8-16] | 9959 |
| **Getting up in the morning** | “Very Easy” | 10687 | “Not at all easy”<br>“Not very easy” | 3554 |
| **Morning/Evening chronotype** | “Definitely a ‘morning’ person” | 7145 | “Definitely an ‘evening’ person” | 2398 |
| **Daytime nap** | “Never/rarely” | 15915 | “Usually” | 1565 |
| **Daytime sleepiness\*** | “Never/rarely” | 21406 | “Sometimes”<br>“Often” | 6458 |
| **Snoring\*** | “Yes” | 16439 | “No” | 9469 |

### Processing of Imaging data

#### Grey Matter Volume (GMV)

T1-weighted pre-processed images were retrieved from UKB with subsequent computations of voxel-based morphometry (CAT 12.7 (default settings); MNI152 space; 1.5mm isotropic)^44^. For each region of interest (ROI), we computed the GMV using the winsorized mean (limits 10%) of the voxel-wise values using the cortical Schaefer atlas (1000 regions of interest, ROIs)^45^, the Melbourne subcortical atlas (S4 3T, 54 ROIs)^46^, and the Diedrichsen cerebellar atlas (SUIT space, 34 ROIs)^47^. This resulted in 1088 GMV features extracted.

#### Brain Surface

We used the data processed using FreeSurfer 6.0 as provided by the UKB^i^. This includes gray/white matter contrast, pial surface, white matter surface, white matter thickness, and white matter volume from the 68 ROIs of the Desikan-Kiliany parcellation^48^, totaling 328 features.

#### Resting-state Functional Magnetic Resonance Imaging (rsfMRI)

The fractional amplitude of low-frequency fluctuations (fALFF) represents the relative measure of blood oxygenation level-dependent (BOLD) magnetic resonance signal power within the low-frequency band of interest (0.008 - 0.09 Hz, reflecting the spontaneous neural activity of the brain) as compared to the BOLD signal power over the entire frequency spectrum^49^. The LCOR (“local correlation”) is a metric that represents the local coherence for each voxel. It is computed as the average of correlation coefficients between a voxel and a region of neighboring voxels, defined by a 25 mm Gaussian kernel^50^. On the other hand, the GCOR (“global correlation”) represents the node centrality of each voxel and is computed as the average of the correlation coefficients between a voxel and all voxels of the whole brain. These metrics were calculated using MatLab2020b, SPM12^51^, FSL (version 5.0)^52^, and the CONN toolbox^53^. The voxel-wise data was then aggregated parcel-wise by averaging according to the parcellation of the GMV data (see above), resulting in 1087 features for each metric (fALFF, LCOR, and GCOR), totaling 3261 features derived from rsfMRI. Note that Diedrichsen cerebellar atlas produced 33 features for the fMRI data as for some ROIs, there were not enough voxels to compute the values correctly. The number of variables and samples for each neuroimaging feature is described in Supplementary Table 1.

### ML models

In order to evaluate a broad spectrum of possible interactions between features and relations to the targets, we selected five machine learning algorithms, including parametric and non-parametric models, testing for linear and nonlinear relations. We tested a Random Forest^54^, Extremely Randomized Trees (Extra Trees)^55^, Support Vector Machine (SVM)^56^, Logistic Regression (logit), and Stacked Generalization^57^, with different hyperparameter settings, resulting in seven models. Table 2 summarizes the models, including the hyperparameters tested, except for the Stacked Generalization model, which is described below. When more than one hyperparameter value was listed, the best hyperparameter value was selected using nested cross-validation (CV), using a grid search approach with a stratified 5-fold CV. The Stacked Generalization model consisted of a Linear SVM with heuristic C^58^ (model LinearSVMHC) for each type of neuroimaging feature (GMV, Surface, fALFF, GCOR, and LCOR) as the first level. The output of each of these five models were used as features of a second-level logistic regression model. For training the second-level model, the out-of-sample predictions of the first-level models were obtained using a stratified 5-fold CV scheme. An overview of the general methodological approach from brain-images and questionaires data to the evaluation of ML models is depicted in Figure 1.

**Figure 1.**
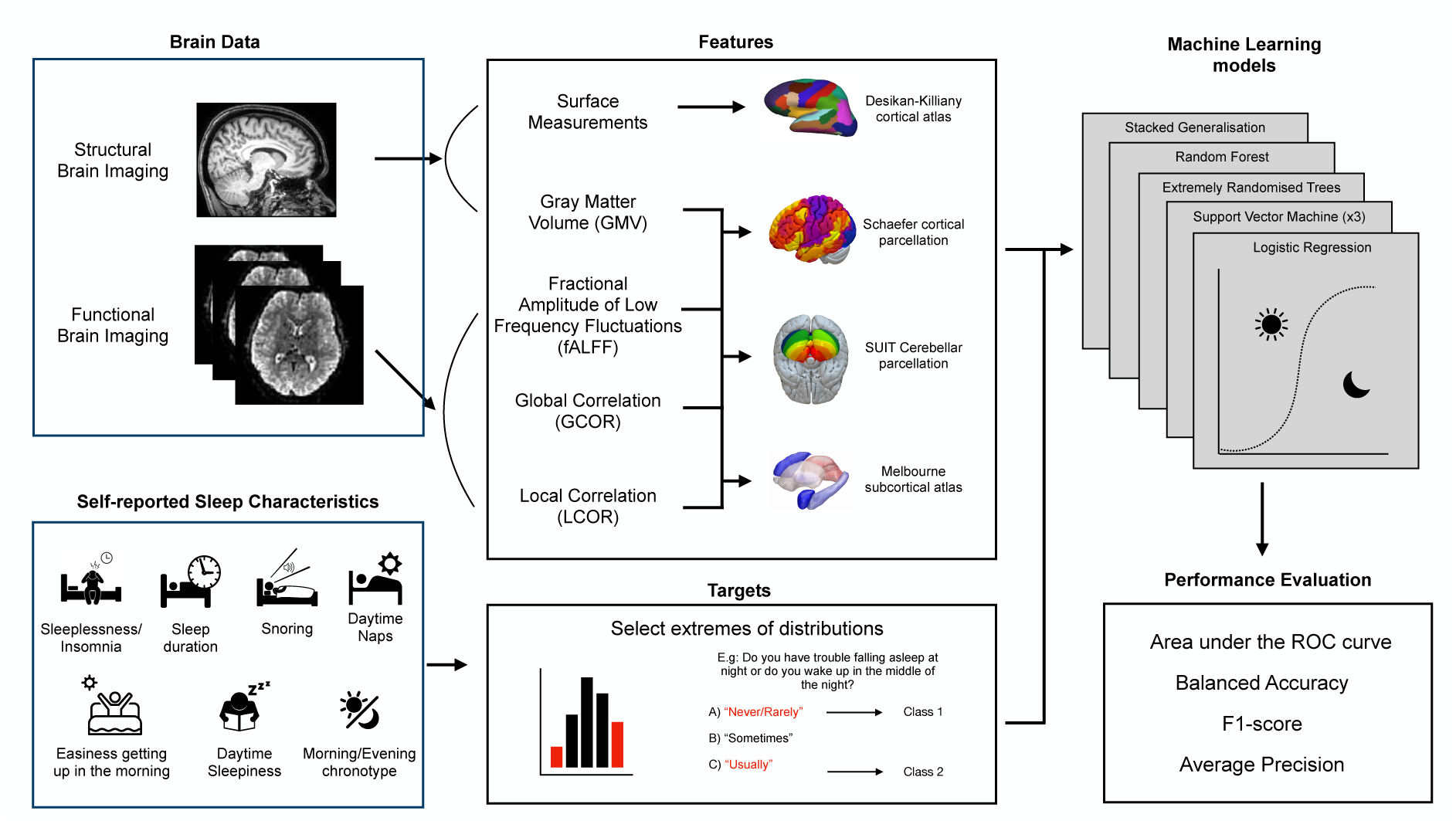
Overview of the methodology. The brain images were processed in order to obtain cortical and subcortical features, both from structural and functional brain imaging. Answers for the UKB questionnaire were binarized by selecting the extremes of the distributions as described in Table 1. We then evaluated the out-of-sample performance of 7 different ML- models, independently for each SH-related characteristic.

**Table 2:** List of models tested, including learning algorithms and hyperparameters evaluated.

| # | Name | Learning Algorithm | Hyperparameter | Values |
| --- | --- | --- | --- | --- |
| 1 | GSET | Extra Trees | estimators | 200, 500 |
|  |  |  | Criterion | Gini, entropy, log loss |
|  |  |  | Max features | Sqrt, log2 |
| 2 | GSRF | Random Forest | estimators | 200, 500 |
|  |  |  | Criterion | Gini, entropy, log loss |
|  |  |  | Max features | Sqrt, log2 |
| **3** | GSSVM-RBF | SVM | Kernel | Rbf |
| | | | C | $1e^{-4}, 1e^{-3}, 1e^{-2}, 1e^{-1}, 1, 10, 100, 1e^4, 1e^5, 1e^6$ |
| | | | Gamma | $1e^{-7}, 1e^{-6}, 1e^{-5}, 1e^{-4}, 1e^{-3}, 1e^{-2}, 1e^{-1}, 1, 10, 100, 1e^4$ |
| **4** | GSLinearSVM | Linear SVM | C | $1e^{-4}, 1e^{-3}, 1e^{-2}, 1e^{-1}, 1, 10, 100, 1e^4, 1e^5, 1e^6$ |
| **5** | LinearSVMHC | Linear SVM | C | heuristic<sup>58</sup> |
|  |  |  | dual | False |
|  |  |  | penalty | L1 |
| **6** | LogitHC | Logit | C | heuristic |
|  |  |  | dual | False |
|  |  |  | penalty | L1 |

### Model evaluation

The available data was first split into 70% training and 30% hold-out test sets to avoid data leakage. Then, the generalization performance of the models (i.e. the capacity to generalize to unseen data) was evaluated on the training set using a stratified 5-fold cross-validation scheme, repeated five times, resulting in 25 evaluation runs. Finally, to validate the CV performance estimation, the models were retrained on the full training set and tested on the hold-out test set. To evaluate different aspects of model performance, such as the trade-off between specificity and sensitivity, we computed two threshold-dependent metrics, namely balanced accuracy and F1 score and two threshold-independent metrics, area under the receiver-operator characteristic (ROC) curve and average precision. Balanced accuracy is computed as the relative number of correct predictions over the total samples, weighted by the number of elements in each class so that the chance level is set at 0.5 and 1 would mean a perfect classification. The F1 score is the harmonic mean between precision and recall^59^. In short, it measures the model’s balanced ability to detect positives (recall = sensitivity) and to have high precision (= positive predictive value), that is, a low rate of false-positive detections. The area under the receiver operating characteristic (ROC) curve (ROC-AUC) provides an aggregate measure of performance across all possible classification thresholds by plotting the true-positive rate (sensitivity) over the false-positive rate (1 – specificity) for each threshold level. Shortly, ROC-AUC can be interpreted as the probability that given two predictions, the model ranks them in the correct order. A perfect model with sensitivity and specificity being equal to 1 at all threshold levels, will have a ROC-AUC of 1, while random guessing will result in ROC-AUC of 0.5^59^. Given that ROC-AUC is skewed for imbalanced datasets, which is the case for all the SH dimensions (see Table 1), a more suitable metric is the area under the Precision-Recall curve^60^, also known as average precision. This metric considers both recall and precision like the F1-score, but across all thresholds as the ROC-AUC does. A perfect model will yield an average precision of 1, while chance levels depend on class balance.

To obtain reference values for each metric, we used the performance of two baseline models, which do not use the features but rely solely on the distribution of classes during training time. A first baseline model named *majority* always predicts the value of the most frequent class in the training set. A second baseline model named *chance* draws random predictions weighted by the number of training samples in each class. All models for each SH dimension were evaluated using the same 5 x 5 CV folds. We then used the corrected paired Student’s t-test for comparing the CV performance of the machine learning models^61^ and corrected for multiple comparisons (across models) using the Bonferroni method. All the analysis described was implemented using Julearn^62^ and Scikit-learn^63^. The codes are available on GitHub: https://github.com/juaml/ukb_sleep_prediction.

### Testing for Confounding bias

To verify that the obtained results were strictly related to brain structure and function, irrespectively of demographic variables such as age and sex, we employed the partial and full confounder statistical tests^64^. This test, developed following the conditional independence testing framework^65^, uses permutation testing to evaluate the independence between pairs of variables, given a potentially high-dimensional random variable that may contain confounding factors. The partial confounder test is used to evaluate if there is a partial confounder bias in the predictions. The null hypothesis states that there is no confounder bias in the data given the target variable (i.e. predictions are independent from the confounder). If the p-value of the partial test is below the threshold (p<0.05), then the null hypothesis can be rejected, indicating that there is an association between the predictions and the confounder. On the other hand, the full confounder test evaluates the null hypothesis that model is entirely driven by the confound. A p-value below the threshold (p<0.05) indicates the model is not fully driven by the confounder. Both tests were parametrized with 1000 permutations and 50 steps for the Markov-chain Monte Carlo sampling.

## Results

We first trained and evaluated all seven models for each of the seven SH-related characteristics, a procedure that took 12.24 core-years, which is approximately 1.5 years on an 8-core desktop computer. For each SH-related characteristic and metric, we selected the best model among the seven competing models according to the performance of the respective metric upon evaluation on 5 x 5 = 25 CV-folds. This resulted in one model per SH-related characteristic and metric, which were then applied to the 30% hold-out test set. The performance of the best model for each SH-related characteristic and metric can be seen in Figure 2. A complete description of the estimated performances for each metric can be seen in Supplementary Table 2.

**Figure 2.**
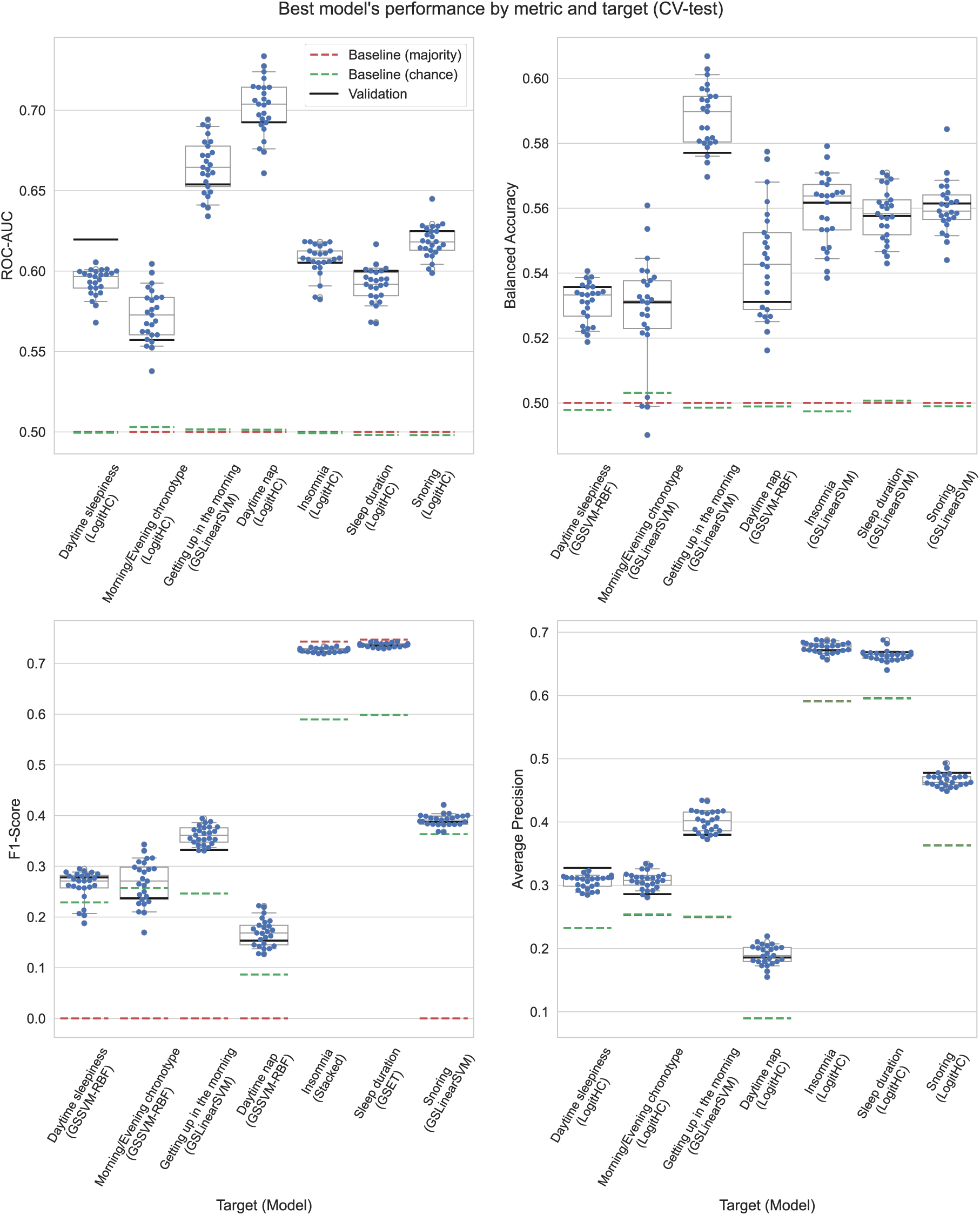
Performances of the best model for each SH-related characteristic. Each blue dot represents the performance obtained at each of the 25 test folds within cross-validation (CV). Boxplots summarize the medians and 95% CI for the underlying distribution. As a reference, dashed red lines depict the mean performance of a model that constantly predicts the most frequent class, green lines depict the mean performance of a model that draws random predictions weighted by the number of samples in each class, and black lines indicate the performance on the hold-out (validation) data.

When only considering the CV performance (which is commonly reported in research settings), some of the SH-related characteristics showed a modest predictability on several metrics. For instance, the best models for *insomnia* and *sleep duration* showed modest balanced accuracy (0.588 and 0.584) and AUC-ROC (0.549 and 0.553) and relatively high F1- score (0.725 and 0.739) and average precision (0.664 and 0.658). However, since some SH-related characteristic have imbalanced classes, it is important to note the performance of the baseline models. For example, the F1 score for insomnia and sleep duration is below the performance of the *majority baseline* model, meaning that a model that simply assigns the majority class to each sample showed a better F1 score. This is, indeed, due to the nature of the F1 scoring, in which chance-level depends on the ratio between classes. The limitation of AUC-ROC with imbalanced data also becomes clear for the *easiness getting up* characteristic, which showed a relatively high AUC-ROC but relatively lower average precision. Furthermore, as cross-validated performances could be overestimated^66^, we evaluated the models on the hold-out data (30% of the samples). The obtained results fall within the confidence intervals of the CV-estimated performances (black lines in Figure 1), suggesting that no over-estimation happened in our case. For more details on the values obtained for each model and SH-related characteristic, see Supplementary Table 3. Overall, our results indicate a weak predictive power but systematically above baseline models for each of the seven SH-related characteristics.

A common ML pitfall with a lack of predictive power is *overfitting*. This occurs when the model closely learns the idiosyncrasies of the training data, thus being incapable of making correct predictions on new unseen samples. To verify that this is not the case, we computed the same metrics for each model but on the training samples. That is, how well each model memorized the training data. The results indicate that while some models were indeed overfitted, at least one model per SH-related characteristic was not (Supplementary Table 4). Given the comparable out-of-sample performance across models for each SH-related characteristic, and that the hyperparameters were selected in nested CV to prevent overfitting, we can safely conclude that overfitting is not a major issue in our results.

We then aimed to identify if the obtained results were purely brain-based predictions or the consequence of confounding bias. In other terms, evaluate if it is possible that the results are simply driven by variables such as the age and sex of the participants, which are known to affect brain structure and function. We employed two different statistical tests: the partial and full confounder tests^64^. The partial confounder test results indicated that among the best models, all of them were partially driven by the age and sex (p<1e-3), except for the models predicting the Morning/Evening chronotype which indicated that there is no evidence to claim that models are partially driven by confounds (p>0.05). On the other hand, the full confounder test indicated that none of the models are fully driven by age and sex (p<=0.001). The full list of p-values for each of the evaluated models can be seen in Supplementary Table 5.

To assess the extent to which age and sex influence the predictions, we trained the learning algorithms from the best models using only these two variables as features. The results, obtained after 6.86 core-years, are depicted in Figure 3. In short, the age and sex models perform similarly for most of the SH-related characteristics, with the exceptions of Sleep Duration, whose prediction by sex and age was worse than the prediction by brain features, and ‘Easiness Getting up in the Morning’ and ‘Daytime Nap’ for which the inverse was true. It is important to note that these models were not optimized for age and sex, but uses the same hyper parametrization as for brain features to serve as a comparison. The full panel including the threshold-dependent metrics are depicted in Supplementary Figure 1.

**Figure 3.**
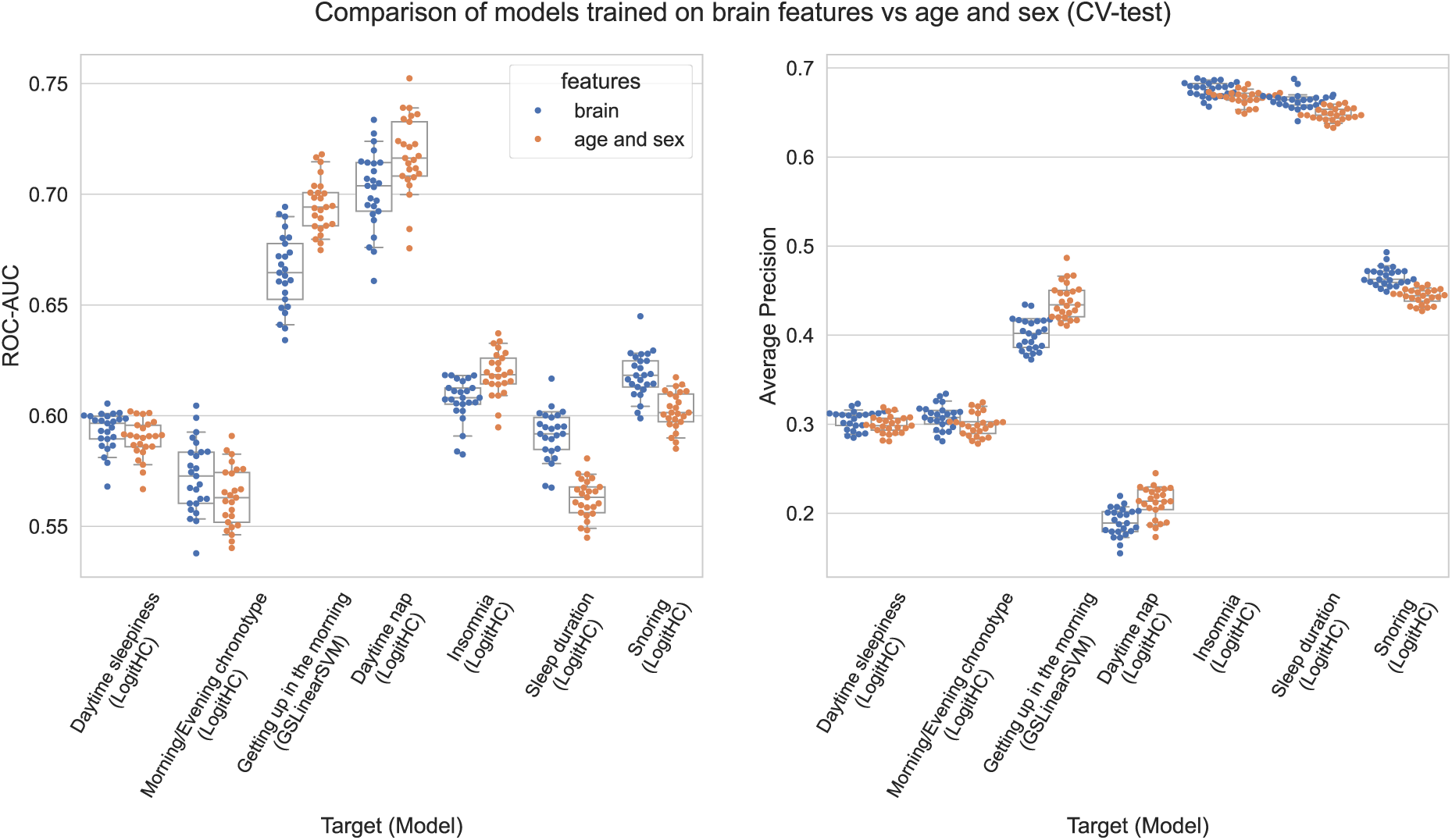
Side-by-side comparison of the best model for each SH-related characteristic using either brain features (blue) or age and sex (orange) indicate that simple demographic variables have the same predictive capacity as complex neuroimaging data in the case of SH-related characteristics. Each dot represents the performance obtained at each of the 25 test folds within cross-validation (CV). Boxplots summarize the medians and 95% CI for the underlying distribution.

## Discussion

The current large-scale study systematically evaluated ML-based predictive analysis for classifying extremes of seven different SH-related characteristics based on multivariate neuroimaging markers in UKB. We covered a large space of multimodal neuroimaging features covering brain structure and function, several ML algorithms in a nested cross-validation setting, and a hold-out test set evaluated on four metrics. Our striking findings demonstrated that the balanced accuracy for predicting SH-related characteristics did not exceed 56%, which indicates that brain structure and function measures could not accurately predict any of the seven SH-related characteristics. The slight improvement over baseline models across the evaluation metrics suggests that the ML algorithms indeed captured some underlying patterns in the data. However, we do not consider these results as high predictive accuracy compared to other brain-imaging-based predictions, such as sex^67,68^, neurodegenerative diseases^69^, and depressive symptoms severity^39^. Furthermore, the comparable predictive performance observed in models trained on age and sex might be why brain-based models can predict above-chance levels. Put differently, we did not observe sufficiently strong evidence to claim that the brain measures can predict SH-related characteristics independently from age and sex. In the following, we discuss the potential reasons for the poor efficacy of multimodal brain features in predicting SH-related characteristics.

### Target issues: Sleep health is a heterogenous concept

Our findings align with previous large-scale sample studies using e.g., UKB and ENIGMA-Sleep datasets that did not observe an association between brain structure and insomnia symptoms^8,23^ and sleep duration^4^. SH has a heterogeneous definition across different general population datasets, as well as clinical samples. Although some studies used a standard sleep questionnaire such as the Pittsburgh Sleep Quality Index (PSQI) to assess sleep quality or the Regulatory Satisfaction Alertness Timing Efficiency Duration (RU-SATED) questionnaire as a valid measure of SH^70^, the UKB did not use those standard questionnaires. Instead, seven self-reported questions were provided about sleep duration, difficulties in getting up in the morning, chronotype, nap, daytime sleepiness, and two measures of clinical conditions such as insomnia symptoms and snoring. Considering these single questions for various SH domains could have affected the clarity and meaningfulness of the measured SH characteristics. Furthermore, the accuracy of self-report sleep assessment based on seven single items and selective participation biases to answer those questions could have led to measurement issues, which have been highlighted previously ^71^.

Another critical aspect is differentiating the sleep-related symptoms of insomnia and snoring in the general population from clinical conditions. It is well-documented that Insomnia disorder is a heterogeneous condition with different subtypes with noticeable inconsistencies in terms of pathophysiology, symptomatology, and treatment response^8,17,25,72–75^. According to the third edition of the International Classification of Sleep Disorders (ICSD-3)^76^, significant daytime dysfunction and having adequate opportunity and circumstances to sleep are essential diagnostic criteria for insomnia disorder. Similarly, snoring can have several etiologies beyond it being a cardinal symptom of OSA, including genetic factors, obesity, nasal blockages, alcohol abuse, smoking, or medications^77^. Thus, relying on a single question about sleep problems is not sufficient to define clinical insomnia disorder or OSA.

Additionally, the imbalance in target labels influences model performance, hindering the learning of sufficient information for accurate classification. Particularly for SH-related characteristics such as ‘Easiness Getting up in the Morning,’ ‘Day Naps,’ and ‘Daytime Dozing,’ the uneven distribution of target labels have resulted in models achieving moderate ROC-AUC scores around 0.6 while the balanced accuracy remained at the chance level of approximately 0.5. This discrepancy between ROC-AUC and balanced accuracy highlights the challenges in achieving fairness and robustness in the models’ predictive capabilities when dealing with imbalanced target datasets. An imbalanced target affects both the learning and interpretation of threshold-dependent metrics^78^. Thus, our conclusions regarding the limited predictive capacity are based on the ROC-AUC and Average Precision metrics, which are threshold-independent, and have been suggested to be preferable for drawing scientific conclusions^60,79^.

The SH-related characteristics in the UKB sample do not represent cross-country sleep differences well. Recently, data from 63 countries showed that individuals from East Asia tend to sleep less and participants from East Europe report longer sleep duration^80^. Similarly, another study on ∼220,000 wearable device users in 35 Countries observed shorter sleep duration, later sleep timing, and less sleep efficiency in East Asia compared with Western Europe, North America, and Oceania, probably due to social- and work-related cultural differences regarding the coping with inadequate sleep and sleep debt ^81^. Moreover, there are significant differences in daytime napping across cultures, being more common in non-Western countries^81^. Of note, however, 10% of the UKB participants reported regular daily naps (Table 1).

### Input features issues: regional brain measurements

Our results also suggest that the neuroimaging features applied in our study may not capture the full spectrum of brain-related features relevant to SH or that the selected features may not be sensitive enough to the subtleties of SH. Moreover, it raises the possibility that current feature sets are insufficiently granular to mirror the complex biological underpinnings of SH. The low performance of the models in predicting SH dimensions, therefore, points to the need for a deeper investigation into more sensitive and comprehensive neuroimaging metrics that can better encapsulate the factors influencing SH. SH might be associated with brain circuits that can be captured, e.g., via seed-based structural or FC measures rather than local brain abnormalities that we used from brain parcels, including GMV, gray/white matter contrast, pial surface, white matter surface, white matter thickness, and white matter volume, LCOR, and fALFF. It has been reported that insomnia symptoms were associated with higher FC within the DMN and FPN and lower FC between the DMN and SN^17^. Wang and colleagues also found that SH dimensions are correlated with disrupted FC patterns in the attentional and thalamic networks in several datasets^7^. Another study using UKB data found associations between SH and FC and structural connectivity. Within-network hyperconnectivity in DMN, FPN, and SN has been observed in healthy subjects and patients with mild cognitive impairment with insomnia symptoms, while patients with Alzheimer’s disease and insomnia symptoms showed hypoconnectivity in those networks^16^. Although we included GCOR, representing functional correlations between a given voxel and other brain voxels (i.e., degree centrality), it didn’t improve the prediction when used as an input with local markers together. Recently, Lynch et al. performed 62 repeated neuroimaging measurements in major depressive disorder (MDD). Using precision functional mapping, they identified FC changes in the frontostriatal circuits that predicted future depressive symptoms^82^ Thus, future studies could explore network-based and white matter integrity metrics as input features or longitudinal precision functional mapping to predict SH in UKB.

Our results also remind us to think beyond the brain feature modalities. Recently, we observed that sleep quality and anxiety robustly predict depressive symptom severity across three independent datasets. Still, brain structural and functional features could not predict depressive symptoms, which indicated that parcellated brain imaging data may not be beneficial in predicting mental health^83^. A large-scale study by the ENIGMA-Anxiety Consortium utilized ML to analyze neuroanatomical data for youth anxiety disorders, and also achieved only modest classification accuracy (AUC 0.59–0.63)^84^. This parallels findings from extensive ML optimization efforts with MDD, which observed mean accuracies in distinguishing patients from controls that ranged from 48.1% to 62.0% only, even when additionally provided with polygenic risk scores,, casting doubt on the potential diagnostic relevance of neuroimaging and genetic biomarkers for MDD^85^. Similarly, the ENIGMA-MDD consortium’s multi-site study^86^ achieved a balanced accuracy of only about 62% in classifying MDD versus healthy controls, which further dropped to approximately 52% after harmonization for site effect. Random chance accuracy was also observed across various stratified groups. These findings may point to an alternative view that complex psychiatric conditions such as sleep disturbance or depression represent deficits in the brain-body interaction, which suggests that body organ health measurements, such as metabolic and cardiovascular systems, in addition to brain imaging, should be considered^87,88^.

### ML-related issues

Following proper ML pipelining practices such as nested CV and grid search for meticulous hyperparameter tuning—methods that typically enhance a model’s capacity to generalize— our models did not achieve high predictive performance. Our study’s low classification performance highlights the inherent challenges in developing models that accurately capture the complex nature of SH using brain imaging data. Machine learning models are designed to discern patterns and generalize findings to new, unseen data. However, like any statistical analysis, ML is challenged when the target labels are unreliable^89^. We reduced the uncertainty in the labeling to some degree by using extreme values for each SH-related characteristic. This should make learning easier for the ML algorithms and boost accuracy. The low performance observed despite this simplification suggests that the prediction of SH-related characteristics as a continuum could be more challenging. Difficulty in creating generalizable ML models arises from potential heterogeneity in how SH is reflected in the brain. In this case, the ML models will not be able to learn a consistent pattern, leading to low performance. Further analysis of SH subtypes and more refined scales are needed to discern this possibility. Finally, several of our classification tasks were imbalanced, i.e. one of the classes was much more frequently present than the other. Such imbalance can lead to biased ML models, which in turn lack generalization ability. To this end, we employed AUC-ROC and average precision metrics to evaluate the ML pipelines. These metrics are independent of a threshold used for dichotomization and thus suitable for characterizing the performance in imbalanced datasets, particularly with tree ensemble models^78^.

### Strengths, limitations, and future directions

The present study has several advantages over other case-control SH-brain studies. Here, we calculated 4677 structural and functional brain features as input features from 28,088 participants from the UKB and applied several ML algorithms to classify the extremes of seven SH-related characteristics. In particular, 1) including diverse and multimodal neuroimaging metrics is crucial. Multimodal data enriches the ML analysis, allowing for a more comprehensive exploration and interpretation of the neurobiological correlates of SH at both structural and functional levels; 2) we leveraged the detailed features provided by the Schaefer atlas (1000 ROIs), which is supported by our ample sample size. This approach assumes that if relevant information is present in an ROI, our models—given their complexity—are equipped to detect it, whether the information is concentrated within a single ROI or dispersed across several regions; 3) we carefully designed our ML analyses using fully separated train and test samples to avoid any leakage of the test set into the model, which is a common oversight in some ML studies^90^; 4) the ML analyses were conducted using several rather different algorithms including Random Forest, Extremely Randomized Trees, Support Vector Machine, Logistic Regression, and Stacked generalization; 5) we applied a grid search-based hyperparameter optimization to prevent overfitting and increase the generalizability of our findings.

Our results should be interpreted within the context of the study’s limitations and the nascent state of this field. This study is based on seven proband answers to SH-related questions and did not include any objective sleep assessment such as polysomnography. Although polysomnography is recommended as a gold-standard objective measure for diagnosing several sleep disorders, including obstructive sleep apnea, its validity for insomnia or sleep quality assessment remains disputed^91^. Moreover, some evidence showed only a weak association between the subjective sleep measurement (e.g., PSQI) and polysomnography in patients with insomnia disorder^92^. Here, we focused on self-reported information on SH. Thus, future studies should consider performing an ML analysis of objective sleep data and comparing it with the analysis of subjective data. Future studies could apply normative modelling, a technique that studies deviations from population norms to show the range of inter-individual differences in brain structure. Unlike traditional case-control paradigms that rely on common neurobiological factors across all subjects, normative modelling focuses on individual deviations from normal patterns, making it a promising approach to consider inter-individual variability in brain expression of SH ^93,94^. Furthermore, longitudinal studies can help identify the long-term interaction between the SH and the brain together with well-characterized sleep measurements from collaborative research groups, e.g., the ENIGMA-Sleep consortium ^95^, to provide replicable results.

## Conclusion

The present extensive ML study using a large population sample demonstrated that multimodal neuroimaging markers had low efficacy in separating the extremes of various sleep health-related characteristics UKB. This suggests that the interaction between sleep health and brain organization may be more complex to be captured with the current ML models and neuroimaging features. While our methodological approach is comprehensive and aims to establish links between neuroimaging features and SH dimensions, this study acknowledges the complexity of interpreting neuroimaging in the context of sleep health. We need future cross-sectional and longitudinal studies considering brain circuits, objective sleep measurements, and cross-country sleep assessments to evaluate the sophisticated brain-sleep interplay.

## Acknowledgements

This project was developed under funding from the Helmholtz Imagining grants NimRLS (ZT-I-PF-4-010) and BrainShapes (ZT-I-PF-4-062). This research has been conducted using data from UK Biobank resources (application number 41655). All data used in this study are publicly accessible from UK Biobank via their standard data access procedure (http://www.ukbiobank.ac.uk/).

## Footnotes

i see https://git.fmrib.ox.ac.uk/falmagro/UK_biobank_pipeline_v_1/-/tree/master/bb_FS_pipeline for the exact pipeline used.

## References

1. Walker, M. P. Sleep essentialism. Brain 144, 697–699 (2021).

2. Cheng, W., Rolls, E. T., Ruan, H. & Feng, J. Functional Connectivities in the Brain That Mediate the Association Between Depressive Problems and Sleep Quality. JAMA Psychiatry 75, 1052 (2018).

3. Ell, J. et al. Sleep health dimensions and shift work as longitudinal predictors of cognitive performance in the UK Biobank cohort. SLEEP 46, zsad093 (2023).

4. Fjell, A. M. et al. No phenotypic or genotypic evidence for a link between sleep duration and brain atrophy. Nat Hum Behav (2023) doi:10.1038/s41562-023-01707-5.

5. Li, Y. et al. The brain structure and genetic mechanisms underlying the nonlinear association between sleep duration, cognition and mental health. Nat Aging 2, 425–437 (2022).

6. Tahmasian, M. et al. The interrelation of sleep and mental and physical health is anchored in grey-matter neuroanatomy and under genetic control. Commun Biol 3, 1–13 (2020).

7. Wang, Y. et al. Covariance patterns between sleep health domains and distributed intrinsic functional connectivity. Nat Commun 14, 7133 (2023).

8. Weihs, A. et al. Lack of structural brain alterations associated with insomnia: findings from the ENIGMA-Sleep Working Group. Journal of Sleep Research 32, e13884 (2023).

9. Buysse, D. J. Sleep Health: Can We Define It? Does It Matter? Sleep 37, 9–17 (2014).

10. Bycroft, C. et al. The UK Biobank resource with deep phenotyping and genomic data. Nature 562, 203–209 (2018).

11. Miller, K. L. et al. Multimodal population brain imaging in the UK Biobank prospective epidemiological study. Nat Neurosci 19, 1523–1536 (2016).

12. Arora, N. et al. Self-reported insomnia symptoms, sleep duration, chronotype and the risk of acute myocardial infarction (AMI): a prospective study in the UK Biobank and the HUNT Study. Eur J Epidemiol 38, 643–656 (2023).

13. Cribb, L. et al. Sleep Regularity and Mortality: A Prospective Analysis in the UK Biobank. 2023.04.14.23288550 Preprint at 10.1101/2023.04.14.23288550 (2023).

14. Kyle, S. D. et al. Sleep and cognitive performance: cross-sectional associations in the UK Biobank. Sleep Medicine 38, 85–91 (2017).

15. Omidvarnia, A. et al. Is resting state fMRI better than individual characteristics at predicting cognition? 2023.02.18.529076 Preprint at 10.1101/2023.02.18.529076 (2023).

16. Elberse, J. D. et al. The interplay between insomnia symptoms and Alzheimer’s disease across three main brain networks. Sleep zsae145 (2024) doi:10.1093/sleep/zsae145.

17. Holub, F. et al. Associations between insomnia symptoms and functional connectivity in the UK Biobank cohort (n = 29,423). Journal of Sleep Research 32, e13790 (2023).

18. Mohajer, B. et al. Gray matter volume and estimated brain age gap are not linked with sleep-disordered breathing. Human Brain Mapping 41, 3034–3044 (2020).

19. Akradi, M. et al. How is sleep-disordered breathing linked with biomarkers of Alzheimer’s disease? 2023.08.16.23294054 Preprint at 10.1101/2023.08.16.23294054 (2023).

20. André, C. et al. Association of Sleep-Disordered Breathing With Alzheimer Disease Biomarkers in Community-Dwelling Older Adults: A Secondary Analysis of a Randomized Clinical Trial. JAMA Neurology 77, 716–724 (2020).

21. Tahmasian, M. et al. Structural and functional neural adaptations in obstructive sleep apnea: An activation likelihood estimation meta-analysis. Neuroscience & Biobehavioral Reviews 65, 142–156 (2016).

22. González, K. A. et al. Sleep duration and brain MRI measures: Results from the SOL-INCA MRI study. Alzheimer’s & Dementia 20, 641–651 (2024).

23. Schiel, J. E. et al. Associations between sleep health and grey matter volume in the UK Biobank cohort (*n* = 33 356). Brain Communications 5, fcad200 (2023).

24. Stolicyn, A. et al. Comprehensive assessment of sleep duration, insomnia, and brain structure within the UK Biobank cohort. Sleep zsad274 (2023) doi:10.1093/sleep/zsad274.

25. Schiel, J. E. et al. Associations Between Sleep Health and Amygdala Reactivity to Negative Facial Expressions in the UK Biobank Cohort. Biological Psychiatry 92, 693–700 (2022).

26. Tsiknia, A. A., Parada, H., Banks, S. J. & Reas, E. T. Sleep quality and sleep duration predict brain microstructure among community-dwelling older adults. Neurobiology of Aging 125, 90–97 (2023).

27. Tai, X. Y., Chen, C., Manohar, S. & Husain, M. Impact of sleep duration on executive function and brain structure. Commun Biol 5, 201 (2022).

28. Fjell, A. M. et al. Is Short Sleep Bad for the Brain? Brain Structure and Cognitive Function in Short Sleepers. J. Neurosci. 43, 5241–5250 (2023).

29. Norbury, R. Diurnal Preference and Grey Matter Volume in a Large Population of Older Adults: Data from the UK Biobank. 18, 3 (2020).

30. Williams, J. A. et al. Genetically mediated associations between chronotype and neuroimaging phenotypes in the UK Biobank: a Mendelian randomisation study. 2023.08.31.555801 Preprint at 10.1101/2023.08.31.555801 (2023).

31. Baril, A.-A. et al. Self-reported sleepiness associates with greater brain and cortical volume and lower prevalence of ischemic covert brain infarcts in a community sample. Sleep 45, zsac185 (2022).

32. Kendler, K. S. Toward a Philosophical Structure for Psychiatry. AJP 162, 433–440 (2005).

33. Bzdok, D. & Yeo, B. T. T. Inference in the age of big data: Future perspectives on neuroscience. NeuroImage 155, 549–564 (2017).

34. Woo, C.-W., Chang, L. J., Lindquist, M. A. & Wager, T. D. Building better biomarkers: brain models in translational neuroimaging. Nat Neurosci 20, 365–377 (2017).

35. Varoquaux, G. et al. Assessing and tuning brain decoders: Cross-validation, caveats, and guidelines. NeuroImage 145, 166–179 (2017).

36. Vieira, S., Pinaya, W. H. L. & Mechelli, A. Using deep learning to investigate the neuroimaging correlates of psychiatric and neurological disorders: Methods and applications. Neuroscience & Biobehavioral Reviews 74, 58–75 (2017).

37. Afshani, M. et al. Discriminating Paradoxical and Psychophysiological Insomnia Based on Structural and Functional Brain Images: A Preliminary Machine Learning Study. Brain Sciences 13, 672 (2023).

38. Goldstein-Piekarski, A. N., Holt-Gosselin, B., O’Hora, K. & Williams, L. M. Integrating sleep, neuroimaging, and computational approaches for precision psychiatry. Neuropsychopharmacol. 45, 192–204 (2020).

39. Olfati, M. et al. Prediction of depressive symptoms severity based on sleep quality, anxiety, and gray matter volume: a generalizable machine learning approach across three datasets. eBioMedicine 108, (2024).

40. Murdoch, W. J., Singh, C., Kumbier, K., Abbasi-Asl, R. & Yu, B. Definitions, methods, and applications in interpretable machine learning. Proceedings of the National Academy of Sciences 116, 22071–22080 (2019).

41. Fan, Z. et al. Mapping sleep’s phenotypic and genetic links to the brain and heart: a systematic analysis of multimodal brain and cardiac images in the UK Biobank. 2022.09.08.22279719 Preprint at 10.1101/2022.09.08.22279719 (2022).

42. Alfaro-Almagro, F. et al. Image processing and Quality Control for the first 10,000 brain imaging datasets from UK Biobank. NeuroImage 166, 400–424 (2018).

43. Goodman, M. O. et al. Genome-wide association analysis of composite sleep health scores in 413,904 individuals. 2024.02.02.24302211 Preprint at 10.1101/2024.02.02.24302211 (2024).

44. Gaser, C. et al. CAT – A Computational Anatomy Toolbox for the Analysis of Structural MRI Data. 2022.06.11.495736 Preprint at 10.1101/2022.06.11.495736 (2023).

45. Schaefer, A. et al. Local-Global Parcellation of the Human Cerebral Cortex from Intrinsic Functional Connectivity MRI. Cerebral Cortex 28, 3095–3114 (2018).

46. Tian, Y. E., Margulies, D. S., Breakspear, M. & Zalesky, A. Topographic organization of the human subcortex unveiled with functional connectivity gradients. 2020.01.13.903542 Preprint at 10.1101/2020.01.13.903542 (2020).

47. Diedrichsen, J., Balsters, J. H., Flavell, J., Cussans, E. & Ramnani, N. A probabilistic MR atlas of the human cerebellum. NeuroImage 46, 39–46 (2009).

48. Desikan, R. S. et al. An automated labeling system for subdividing the human cerebral cortex on MRI scans into gyral based regions of interest. NeuroImage 31, 968–980 (2006).

49. Zou, Q.-H. et al. An improved approach to detection of amplitude of low-frequency fluctuation (ALFF) for resting-state fMRI: fractional ALFF. J Neurosci Methods 172, 137– 141 (2008).

50. Deshpande, G., LaConte, S., Peltier, S. & Hu, X. Integrated local correlation: A new measure of local coherence in fMRI data. Human Brain Mapping 30, 13–23 (2009).

51. Friston, K. J., Ashburner, J., Kiebel, S., Nichols, T. & Penny, W. Statistical Parametric Mapping: The Analysis of Functional Brain Images. (Elsevier / Academic Press, Amsterdam Boston, 2007).

52. Jenkinson, M., Beckmann, C. F., Behrens, T. E. J., Woolrich, M. W. & Smith, S. M. FSL. Neuroimage 62, 782–790 (2012).

53. Whitfield-Gabrieli, S. & Nieto-Castanon, A. Conn: a functional connectivity toolbox for correlated and anticorrelated brain networks. Brain Connect 2, 125–141 (2012).

54. Breiman, L. Random forests. Machine Learning 45, 5–32 (2001).

55. Geurts, P., Ernst, D. & Wehenkel, L. Extremely randomized trees. Machine Learning 63, 3– 42 (2006).

56. Cortes, C. & Vapnik, V. Support-vector networks. Mach Learn 20, 273–297 (1995).

57. Wolpert, D. H. Stacked generalization. Neural Networks 5, 241–259 (1992).

58. R: Fast Heuristics For The Estimation Of the C Constant Of A… https://search.r-project.org/CRAN/refmans/LiblineaR/html/heuristicC.html.

59. Hastie, T., Friedman, J. & Tibshirani, R. *The Elements of Statistical Learning*. (Springer New York, New York, NY, 2001). doi:10.1007/978-0-387-21606-5.

60. Davis, J. & Goadrich, M. The relationship between Precision-Recall and ROC curves. in Proceedings of the 23rd international conference on Machine learning 233–240 (Association for Computing Machinery, New York, NY, USA, 2006). doi:10.1145/1143844.1143874.

61. Nadeau, C. & Bengio, Y. Inference for the Generalization Error. Machine Learning 52, 239– 281 (2003).

62. Hamdan, S. et al. Julearn: an easy-to-use library for leakage-free evaluation and inspection of ML models. Gigabyte 2024, 1–16 (2024).

63. Pedregosa, F. et al. Scikit-learn: Machine Learning in Python. Journal of Machine Learning Research 12, 2825–2830 (2012).

64. Spisak, T. Statistical quantification of confounding bias in machine learning models. GigaScience 11, giac082 (2022).

65. Berrett, T. B., Wang, Y., Barber, R. F. & Samworth, R. J. The conditional permutation test for independence while controlling for confounders. Preprint at 10.48550/arXiv.1807.05405 (2019).

66. Varoquaux, G. Cross-validation failure: Small sample sizes lead to large error bars. Neuroimage (2017).

67. Schulz, M.-A., Bzdok, D., Haufe, S., Haynes, J.-D. & Ritter, K. Performance reserves in brain-imaging-based phenotype prediction. Cell Reports 43, 113597 (2024).

68. Wiersch, L. et al. Accurate sex prediction of cisgender and transgender individuals without brain size bias. Sci Rep 13, 13868 (2023).

69. Kasper, J. et al. Local synchronicity in dopamine-rich caudate nucleus influences Huntington’s disease motor phenotype. Brain 146, 3319–3330 (2023).

70. Ravyts, S. G., Dizerzewski, J. M., Perez, E., Donovan, E. K. & Dautovich, N. Sleep Health as Measured by RU SATED: A Psychometric Evaluation. Behav Sleep Med 19, 48–56 (2021).

71. Schoeler, T., Pingault, J.-B. & Kutalik, Z. Self-report inaccuracy in the UK Biobank: Impact on inference and interplay with selective participation. 2023.10.06.23296652 Preprint at 10.1101/2023.10.06.23296652 (2023).

72. Blanken, T. F. et al. Insomnia disorder subtypes derived from life history and traits of affect and personality. The Lancet Psychiatry 6, 151–163 (2019).

73. Bresser, T. et al. Insomnia subtypes have differentiating deviations in brain structural connectivity. Biological Psychiatry (2024) doi:10.1016/j.biopsych.2024.06.014.

74. Emamian, F. et al. Alterations of Subcortical Brain Structures in Paradoxical and Psychophysiological Insomnia Disorder. Frontiers in Psychiatry 12, (2021).

75. Reimann, G. M. et al. Convergent abnormality in the subgenual anterior cingulate cortex in insomnia disorder: A revisited neuroimaging meta-analysis of 39 studies. Sleep Medicine Reviews 71, 101821 (2023).

76. Sateia, M. J. International classification of sleep disorders-third edition: highlights and modifications. Chest 146, 1387–1394 (2014).

77. Campos, A. I. et al. Insights into the aetiology of snoring from observational and genetic investigations in the UK Biobank. Nat Commun 11, 817 (2020).

78. Collell, G., Prelec, D. & Patil, K. R. A simple plug-in bagging ensemble based on threshold-moving for classifying binary and multiclass imbalanced data. Neurocomputing 275, 330– 340 (2018).

79. Provost, F. J., Fawcett, T. & Kohavi, R. The Case against Accuracy Estimation for Comparing Induction Algorithms. in Proceedings of the Fifteenth International Conference on Machine Learning 445–453 (Morgan Kaufmann Publishers Inc., San Francisco, CA, USA, 1998).

80. Coutrot, A. et al. Reported sleep duration reveals segmentation of the adult life-course into three phases. Nat Commun 13, 7697 (2022).

81. Willoughby, A. R., Alikhani, I., Karsikas, M., Chua, X. Y. & Chee, M. W. L. Country differences in nocturnal sleep variability: Observations from a large-scale, long-term sleep wearable study. Sleep Medicine 110, 155–165 (2023).

82. Lynch, C. J. et al. Frontostriatal salience network expansion in individuals in depression. Nature 633, 624–633 (2024).

83. Olfati, M. et al. Prediction of depressive symptoms severity based on sleep quality, anxiety, and brain: a machine learning approach across three cohorts. (2024) doi:10.1101/2023.08.09.23293887.

84. Bruin, W. B. et al. Brain-based classification of youth with anxiety disorders: transdiagnostic examinations within the ENIGMA-Anxiety database using machine learning. Nat. Mental Health 2, 104–118 (2024).

85. Winter, N. R. et al. A Systematic Evaluation of Machine Learning–Based Biomarkers for Major Depressive Disorder. JAMA Psychiatry (2024) doi:10.1001/jamapsychiatry.2023.5083.

86. Belov, V. et al. Multi-site benchmark classification of major depressive disorder using machine learning on cortical and subcortical measures. Scientific Reports 14, 1084 (2024).

87. Kendler, K. S. Are Psychiatric Disorders Brain Diseases?—A New Look at an Old Question. JAMA Psychiatry (2024) doi:10.1001/jamapsychiatry.2024.0036.

88. Tian, Y. E. et al. Evaluation of Brain-Body Health in Individuals With Common Neuropsychiatric Disorders. JAMA Psychiatry 80, 567–576 (2023).

89. Gell, M. et al. The Burden of Reliability: How Measurement Noise Limits Brain-Behaviour Predictions. 2023.02.09.527898 Preprint at 10.1101/2023.02.09.527898 (2024).

90. Sasse, L., et al. On Leakage in Machine Learning Pipelines. Preprint at 10.48550/arXiv.2311.04179 (2024).

91. Frase, L., Nissen, C., Spiegelhalder, K. & Feige, B. The importance and limitations of polysomnography in insomnia disorder—a critical appraisal. Journal of Sleep Research 32, e14036 (2023).

92. Benz, F. et al. How many hours do you sleep? A comparison of subjective and objective sleep duration measures in a sample of insomnia patients and good sleepers. Journal of Sleep Research 32, e13802 (2023).

93. Marquand, A. F., Rezek, I., Buitelaar, J. & Beckmann, C. F. Understanding Heterogeneity in Clinical Cohorts Using Normative Models: Beyond Case-Control Studies. Biological Psychiatry 80, 552–561 (2016).

94. Rutherford, S. et al. The normative modeling framework for computational psychiatry. Nat Protoc 17, 1711–1734 (2022).

95. Tahmasian, M. et al. ENIGMA-Sleep: Challenges, opportunities, and the road map. Journal of Sleep Research 30, e13347 (2021).

