## Supplementary Material for "Can we predict sleep health based on brain features? A large-scale machine learning study using UK Biobank"

| Features | Atlas | # ROIs | # Samples |
|---|---|---|---|
| **GMV** | Schaefer | 1000 | 39316 |
| | Melbourne Subcortical | 54 | 39172 |
| | Diedrichsen | 34 | 38490 |
| **gray/white matter contrast** | Desikan-Kiliany | 68 | 43107 |
| **Pial surface** | Desikan-Kiliany | 64 | 43107 |
| **White matter surface** | Desikan-Kiliany | 66 | 43107 |
| **White matter thickness** | Desikan-Kiliany | 66 | 43107 |
| **White matter volume** | Desikan-Kiliany | 64 | 43107 |
| **fALFF** | Schaefer | 1000 | 29367 |
| | Melbourne Subcortical | 54 | 29367 |
| | Diedrichsen | 33 | 29367 |
| **LCOR** | Schaefer | 1000 | 29367 |
| | Melbourne Subcortical | 54 | 29367 |
| | Diedrichsen | 33 | 29367 |
| **GCOR** | Schaefer | 1000 | 29367 |
| | Melbourne Subcortical | 54 | 29367 |
| | Diedrichsen | 33 | 29367 |

Table S1. List of features extracted from functional and structural brain imaging, indicating the parcellation applied, the number of ROIs extracted, and the number of participants for which the features were successfully extracted. Note: GMV = Gray Matter Volume, fALFF = Fractional Amplitude of Low-Frequency Fluctuations, LCOR = Local Correlation, GCOR = Global Correlation.

| SH Characteristic | Model | ROC AUC | Balanced Accuracy | F1-Score | Average Precision |
|---|---|---|---|---|---|
| **Daytime Sleepiness** | Extra Trees | 0.581 ± 0.009 | 0.5 ± 0.0 | 0.001 ± 0.001 | 0.288 ± 0.008 |
| | Linear SVM | 0.562 ± 0.014 | 0.527 ± 0.007 | 0.23 ± 0.056 | 0.278 ± 0.011 |
| | Random Forest | 0.575 ± 0.007 | 0.501 ± 0.001 | 0.005 ± 0.004 | 0.282 ± 0.006 |
| | SVM (RBF Kernel) | 0.558 ± 0.009 | **0.531 ± 0.006** | **0.262 ± 0.03** | 0.275 ± 0.005 |
| | Linear SVM (Heuristic C) | 0.581 ± 0.007 | 0.514 ± 0.003 | 0.1 ± 0.01 | 0.295 ± 0.01 |
| | Logit (Heuristic C) | **0.593 ± 0.009** | 0.501 ± 0.001 | 0.008 ± 0.003 | **0.305 ± 0.011** |
| | Stacked | 0.511 ± 0.004 | 0.5 ± 0.0 | 0.002 ± 0.003 | 0.24 ± 0.004 |
| **Morning/ Evening chronotype** | Extra Trees | 0.539 ± 0.021 | 0.5 ± 0.002 | 0.004 ± 0.005 | 0.287 ± 0.016 |
| | Linear SVM | 0.557 ± 0.025 | **0.529 ± 0.017** | 0.262 ± 0.022 | 0.299 ± 0.022 |
| | Random Forest | 0.538 ± 0.019 | 0.501 ± 0.002 | 0.01 ± 0.009 | 0.282 ± 0.014 |

| | | | | | |
|---|---|---|---|---|---|
| | SVM (RBF Kernel) | 0.54 ± 0.018 | 0.521 ± 0.016 | **0.268 ± 0.045** | 0.284 ± 0.016 |
| | Linear SVM (Heuristic C) | 0.549 ± 0.019 | 0.511 ± 0.007 | 0.112 ± 0.023 | 0.292 ± 0.015 |
| | Logit (Heuristic C) | **0.572 ± 0.017** | 0.5 ± 0.0 | 0.0 ± 0.0 | **0.308 ± 0.014** |
| | Stacked | 0.512 ± 0.009 | 0.5 ± 0.0 | 0.0 ± 0.001 | 0.26 ± 0.006 |
| **Easiness getting up in the morning** | Extra Trees | 0.631 ± 0.02 | 0.502 ± 0.002 | 0.011 ± 0.007 | 0.36 ± 0.02 |
| | Linear SVM | **0.665 ± 0.018** | **0.588 ± 0.01** | **0.361 ± 0.018** | **0.401 ± 0.018** |
| | Random Forest | 0.619 ± 0.017 | 0.504 ± 0.003 | 0.026 ± 0.009 | 0.344 ± 0.017 |
| | SVM (RBF Kernel) | 0.634 ± 0.02 | 0.57 ± 0.006 | 0.324 ± 0.014 | 0.372 ± 0.017 |
| | Linear SVM (Heuristic C) | 0.653 ± 0.017 | 0.553 ± 0.008 | 0.25 ± 0.019 | 0.383 ± 0.016 |
| | Logit (Heuristic C) | 0.664 ± 0.015 | 0.515 ± 0.005 | 0.074 ± 0.018 | 0.396 ± 0.016 |
| | Stacked | 0.563 ± 0.01 | 0.515 ± 0.005 | 0.076 ± 0.021 | 0.302 ± 0.011 |
| **Daytime nap** | Extra Trees | 0.631 ± 0.02 | 0.502 ± 0.002 | 0.011 ± 0.007 | 0.36 ± 0.02 |
| | Linear SVM | **0.665 ± 0.018** | **0.588 ± 0.01** | **0.361 ± 0.018** | **0.401 ± 0.018** |
| | Random Forest | 0.619 ± 0.017 | 0.504 ± 0.003 | 0.026 ± 0.009 | 0.344 ± 0.017 |
| | SVM (RBF Kernel) | 0.634 ± 0.02 | 0.57 ± 0.006 | 0.324 ± 0.014 | 0.372 ± 0.017 |
| | Linear SVM (Heuristic C) | 0.653 ± 0.017 | 0.553 ± 0.008 | 0.25 ± 0.019 | 0.383 ± 0.016 |
| | Logit (Heuristic C) | 0.664 ± 0.015 | 0.515 ± 0.005 | 0.074 ± 0.018 | 0.396 ± 0.016 |
| | Stacked | 0.563 ± 0.01 | 0.515 ± 0.005 | 0.076 ± 0.021 | 0.302 ± 0.011 |
| **Insomnia** | Extra Trees | 0.59 ± 0.012 | 0.531 ± 0.007 | 0.724 ± 0.005 | 0.661 ± 0.013 |
| | Linear SVM | 0.588 ± 0.012 | **0.56 ± 0.011** | 0.661 ± 0.009 | 0.663 ± 0.01 |
| | Random Forest | 0.59 ± 0.012 | 0.535 ± 0.007 | 0.721 ± 0.007 | 0.662 ± 0.011 |
| | SVM (RBF Kernel) | 0.595 ± 0.016 | 0.551 ± 0.007 | 0.709 ± 0.016 | 0.668 ± 0.015 |
| | Linear SVM (Heuristic C) | 0.578 ± 0.009 | 0.55 ± 0.008 | 0.671 ± 0.006 | 0.655 ± 0.009 |
| | Logit (Heuristic C) | **0.607 ± 0.01** | 0.553 ± 0.008 | 0.717 ± 0.006 | **0.676 ± 0.009** |
| | Stacked | 0.583 ± 0.009 | 0.535 ± 0.006 | **0.727 ± 0.004** | 0.647 ± 0.007 |
| **Sleep duration** | Extra Trees | 0.572 ± 0.014 | 0.531 ± 0.006 | **0.736 ± 0.004** | 0.645 ± 0.011 |
| | Linear SVM | 0.587 ± 0.008 | **0.558 ± 0.008** | 0.672 ± 0.008 | 0.663 ± 0.009 |
| | Random Forest | 0.573 ± 0.014 | 0.532 ± 0.007 | 0.734 ± 0.005 | 0.648 ± 0.012 |
| | SVM (RBF Kernel) | 0.575 ± 0.014 | 0.544 ± 0.006 | 0.682 ± 0.035 | 0.656 ± 0.011 |
| | Linear SVM (Heuristic C) | 0.581 ± 0.008 | 0.551 ± 0.008 | 0.681 ± 0.009 | 0.657 ± 0.01 |
| | Logit (Heuristic C) | **0.591 ± 0.011** | 0.545 ± 0.005 | 0.731 ± 0.006 | **0.664 ± 0.008** |
| | Stacked | 0.57 ± 0.009 | 0.526 ± 0.004 | 0.735 ± 0.005 | 0.641 ± 0.008 |
| **Snoring** | Extra Trees | 0.574 ± 0.012 | 0.503 ± 0.002 | 0.032 ± 0.006 | 0.423 ± 0.011 |
| | Linear SVM | 0.6 ± 0.009 | **0.56 ± 0.007** | **0.391 ± 0.011** | 0.45 ± 0.01 |
| | Random Forest | 0.571 ± 0.012 | 0.505 ± 0.003 | 0.054 ± 0.012 | 0.42 ± 0.013 |
| | SVM (RBF Kernel) | 0.576 ± 0.008 | 0.547 ± 0.007 | 0.378 ± 0.012 | 0.428 ± 0.009 |
| | Linear SVM (Heuristic C) | 0.592 ± 0.008 | 0.551 ± 0.006 | 0.356 ± 0.011 | 0.444 ± 0.011 |
| | Logit (Heuristic C) | **0.618 ± 0.01** | 0.538 ± 0.005 | 0.234 ± 0.01 | **0.466 ± 0.011** |
| | Stacked | 0.578 ± 0.008 | 0.52 ± 0.005 | 0.138 ± 0.018 | 0.425 ± 0.009 |

Table S2: Results of the out-of-sample evaluation (CV) of the 7 models for each SH-related characteristic. Values represent the mean and standard deviation across CV repetitions.

Values in bold represent the best performance for each metric and SH-related characteristic across the 7 models.

| SH Characteristic | Model | ROC AUC | Balanced Accuracy | F1-Score | Average Precision |
|---|---|---|---|---|---|
| **Daytime Sleepiness** | Random Forest | 0.501 | 0.006 | 0.289 | 0.585 |
| | Linear SVM (Heuristic C) | 0.512 | 0.087 | 0.297 | 0.589 |
| | Linear SVM | 0.53 | 0.262 | 0.266 | 0.547 |
| | Logit (Heuristic C) | 0.502 | 0.012 | 0.327 | 0.62 |
| | SVM (RBF Kernel) | 0.536 | 0.278 | 0.274 | 0.559 |
| | Stacked | 0.501 | 0.006 | 0.238 | 0.51 |
| | Extra Trees | 0.499 | 0.0 | 0.258 | 0.524 |
| **Morning/ Evening chronotype** | Random Forest | 0.5 | 0.006 | 0.256 | 0.527 |
| | Linear SVM (Heuristic C) | 0.505 | 0.09 | 0.289 | 0.564 |
| | Linear SVM | 0.531 | 0.232 | 0.291 | 0.571 |
| | Logit (Heuristic C) | 0.5 | 0.0 | 0.286 | 0.557 |
| | SVM (RBF Kernel) | 0.53 | 0.237 | 0.286 | 0.555 |
| | Stacked | 0.5 | 0.0 | 0.253 | 0.515 |
| | Extra Trees | 0.5 | 0.006 | 0.342 | 0.622 |
| **Easiness getting up in the morning** | Random Forest | 0.502 | 0.025 | 0.331 | 0.61 |
| | Linear SVM (Heuristic C) | 0.549 | 0.24 | 0.362 | 0.639 |
| | Linear SVM | 0.577 | 0.333 | 0.38 | 0.654 |
| | Logit (Heuristic C) | 0.518 | 0.089 | 0.39 | 0.663 |
| | SVM (RBF Kernel) | 0.555 | 0.3 | 0.338 | 0.609 |
| | Stacked | 0.515 | 0.08 | 0.296 | 0.561 |
| | Extra Trees | 0.5 | 0.0 | 0.169 | 0.666 |
| **Daytime nap** | Random Forest | 0.501 | 0.004 | 0.154 | 0.643 |
| | Linear SVM (Heuristic C) | 0.5 | 0.0 | 0.163 | 0.668 |
| | Linear SVM | 0.525 | 0.146 | 0.111 | 0.583 |
| | Logit (Heuristic C) | 0.501 | 0.004 | 0.186 | 0.693 |
| | SVM (RBF Kernel) | 0.531 | 0.154 | 0.113 | 0.586 |
| | Stacked | 0.5 | 0.0 | 0.09 | 0.5 |
| | Extra Trees | 0.535 | 0.719 | 0.661 | 0.586 |
| **Insomnia** | Random Forest | 0.542 | 0.717 | 0.653 | 0.587 |
| | Linear SVM (Heuristic C) | 0.545 | 0.659 | 0.655 | 0.58 |
| | Linear SVM | 0.562 | 0.657 | 0.662 | 0.587 |
| | Logit (Heuristic C) | 0.551 | 0.703 | 0.672 | 0.605 |
| | SVM (RBF Kernel) | 0.555 | 0.705 | 0.667 | 0.599 |
| | Stacked | 0.536 | 0.722 | 0.657 | 0.597 |
| | Extra Trees | 0.533 | 0.735 | 0.646 | 0.579 |
| **Sleep duration** | Random Forest | 0.532 | 0.735 | 0.652 | 0.581 |

| | | | | | |
|---|---|---|---|---|---|
| | Linear SVM (Heuristic C) | 0.554 | 0.686 | 0.663 | 0.586 |
| | Linear SVM | 0.558 | 0.679 | 0.672 | 0.596 |
| | Logit (Heuristic C) | 0.542 | 0.728 | 0.669 | 0.6 |
| | SVM (RBF Kernel) | 0.55 | 0.664 | 0.649 | 0.568 |
| | Stacked | 0.539 | 0.742 | 0.651 | 0.589 |
| | Extra Trees | 0.503 | 0.035 | 0.428 | 0.579 |
| **Snoring** | Random Forest | 0.507 | 0.056 | 0.427 | 0.574 |
| | Linear SVM (Heuristic C) | 0.552 | 0.355 | 0.453 | 0.606 |
| | Linear SVM | 0.561 | 0.387 | 0.461 | 0.613 |
| | Logit (Heuristic C) | 0.542 | 0.255 | 0.478 | 0.625 |
| | SVM (RBF Kernel) | 0.551 | 0.375 | 0.444 | 0.595 |
| | Stacked | 0.524 | 0.171 | 0.431 | 0.582 |
| | Random Forest | 0.501 | 0.006 | 0.289 | 0.585 |

Table S3: Results of the validation sample evaluation of the 7 models for each SH-related characteristic.

| SH Characteristic | Model | ROC AUC | Balanced Accuracy | F1-Score | Average Precision |
|---|---|---|---|---|---|
| **Daytime Sleepiness** | Extra Trees | 1.0 ± 0.0 | 1.0 ± 0.0 | 1.0 ± 0.0 | 1.0 ± 0.0 |
| | Linear SVM | 0.871 ± 0.039 | 0.701 ± 0.068 | 0.548 ± 0.136 | 0.71 ± 0.074 |
| | Random Forest | 1.0 ± 0.0 | 1.0 ± 0.0 | 1.0 ± 0.0 | 1.0 ± 0.0 |
| | SVM (RBF Kernel) | 0.958 ± 0.058 | 0.919 ± 0.105 | 0.891 ± 0.145 | 0.923 ± 0.108 |
| | Linear SVM (Heuristic C) | 0.799 ± 0.002 | 0.56 ± 0.003 | 0.222 ± 0.008 | 0.577 ± 0.004 |
| | Logit (Heuristic C) | 0.668 ± 0.002 | 0.502 ± 0.001 | 0.011 ± 0.004 | 0.381 ± 0.004 |
| | Stacked | 0.537 ± 0.003 | 0.502 ± 0.001 | 0.007 ± 0.005 | 0.274 ± 0.004 |
| **Morning/ Evening chronotype** | Extra Trees | 1.0 ± 0.0 | 1.0 ± 0.0 | 1.0 ± 0.0 | 1.0 ± 0.0 |
| | Linear SVM | 0.906 ± 0.048 | 0.786 ± 0.092 | 0.698 ± 0.134 | 0.799 ± 0.101 |
| | Random Forest | 1.0 ± 0.0 | 1.0 ± 0.0 | 1.0 ± 0.0 | 1.0 ± 0.0 |
| | SVM (RBF Kernel) | 0.983 ± 0.023 | 0.931 ± 0.084 | 0.916 ± 0.104 | 0.974 ± 0.035 |
| | Linear SVM (Heuristic C) | 0.858 ± 0.004 | 0.598 ± 0.008 | 0.333 ± 0.02 | 0.702 ± 0.009 |
| | Logit (Heuristic C) | 0.637 ± 0.007 | 0.5 ± 0.0 | 0.0 ± 0.0 | 0.369 ± 0.007 |
| | Stacked | 0.619 ± 0.024 | 0.5 ± 0.0 | 0.001 ± 0.002 | 0.404 ± 0.025 |
| **Easiness getting up in the morning** | Extra Trees | 1.0 ± 0.0 | 1.0 ± 0.0 | 1.0 ± 0.0 | 1.0 ± 0.0 |
| | Linear SVM | 0.874 ± 0.003 | 0.735 ± 0.006 | 0.619 ± 0.009 | 0.724 ± 0.006 |
| | Random Forest | 1.0 ± 0.0 | 1.0 ± 0.0 | 1.0 ± 0.0 | 1.0 ± 0.0 |
| | SVM (RBF Kernel) | 0.989 ± 0.016 | 0.962 ± 0.048 | 0.953 ± 0.061 | 0.98 ± 0.027 |
| | Linear SVM (Heuristic C) | 0.865 ± 0.004 | 0.651 ± 0.007 | 0.467 ± 0.015 | 0.707 ± 0.007 |
| | Logit (Heuristic C) | 0.713 ± 0.005 | 0.522 ± 0.004 | 0.095 ± 0.014 | 0.458 ± 0.006 |
| | Stacked | 0.661 ± 0.01 | 0.54 ± 0.011 | 0.155 ± 0.04 | 0.442 ± 0.012 |
| **Daytime nap** | Extra Trees | 1.0 ± 0.0 | 1.0 ± 0.0 | 1.0 ± 0.0 | 1.0 ± 0.0 |

| | | | | | |
|---|---|---|---|---|---|
| | Linear SVM | 1.0 ± 0.0 | 0.999 ± 0.002 | 0.999 ± 0.002 | 1.0 ± 0.0 |
| | Random Forest | 1.0 ± 0.0 | 1.0 ± 0.0 | 1.0 ± 0.0 | 1.0 ± 0.0 |
| | SVM (RBF Kernel) | 1.0 ± 0.0 | 0.999 ± 0.001 | 0.999 ± 0.001 | 1.0 ± 0.001 |
| | Linear SVM (Heuristic C) | 0.89 ± 0.003 | 0.503 ± 0.002 | 0.014 ± 0.006 | 0.546 ± 0.012 |
| | Logit (Heuristic C) | 0.737 ± 0.005 | 0.5 ± 0.0 | 0.0 ± 0.0 | 0.223 ± 0.007 |
| | Stacked | 0.501 ± 0.003 | 0.5 ± 0.0 | 0.0 ± 0.0 | 0.093 ± 0.004 |
| **Insomnia** | Extra Trees | 1.0 ± 0.0 | 1.0 ± 0.0 | 1.0 ± 0.0 | 1.0 ± 0.0 |
| | Linear SVM | 0.82 ± 0.002 | 0.732 ± 0.003 | 0.796 ± 0.002 | 0.862 ± 0.002 |
| | Random Forest | 1.0 ± 0.0 | 1.0 ± 0.0 | 1.0 ± 0.0 | 1.0 ± 0.0 |
| | SVM (RBF Kernel) | 0.781 ± 0.033 | 0.671 ± 0.047 | 0.792 ± 0.027 | 0.813 ± 0.028 |
| | Linear SVM (Heuristic C) | 0.822 ± 0.002 | 0.722 ± 0.004 | 0.802 ± 0.002 | 0.864 ± 0.003 |
| | Logit (Heuristic C) | 0.666 ± 0.003 | 0.584 ± 0.003 | 0.738 ± 0.002 | 0.728 ± 0.003 |
| | Stacked | 0.736 ± 0.005 | 0.604 ± 0.006 | 0.77 ± 0.002 | 0.762 ± 0.004 |
| **Sleep duration** | Extra Trees | 1.0 ± 0.0 | 1.0 ± 0.0 | 1.0 ± 0.0 | 1.0 ± 0.0 |
| | Linear SVM | 0.816 ± 0.002 | 0.725 ± 0.003 | 0.799 ± 0.002 | 0.859 ± 0.002 |
| | Random Forest | 1.0 ± 0.0 | 1.0 ± 0.0 | 1.0 ± 0.0 | 1.0 ± 0.0 |
| | SVM (RBF Kernel) | 0.901 ± 0.107 | 0.834 ± 0.176 | 0.898 ± 0.105 | 0.913 ± 0.092 |
| | Linear SVM (Heuristic C) | 0.818 ± 0.003 | 0.717 ± 0.004 | 0.805 ± 0.002 | 0.86 ± 0.002 |
| | Logit (Heuristic C) | 0.661 ± 0.004 | 0.572 ± 0.004 | 0.749 ± 0.001 | 0.726 ± 0.004 |
| | Stacked | 0.736 ± 0.005 | 0.59 ± 0.013 | 0.772 ± 0.004 | 0.763 ± 0.004 |
| **Snoring** | Extra Trees | 1.0 ± 0.0 | 1.0 ± 0.0 | 1.0 ± 0.0 | 1.0 ± 0.0 |
| | Linear SVM | 0.786 ± 0.002 | 0.684 ± 0.003 | 0.575 ± 0.005 | 0.682 ± 0.003 |
| | Random Forest | 1.0 ± 0.0 | 1.0 ± 0.0 | 1.0 ± 0.0 | 1.0 ± 0.0 |
| | SVM (RBF Kernel) | 0.884 ± 0.094 | 0.822 ± 0.136 | 0.766 ± 0.182 | 0.835 ± 0.134 |
| | Linear SVM (Heuristic C) | 0.788 ± 0.002 | 0.671 ± 0.003 | 0.545 ± 0.006 | 0.686 ± 0.004 |
| | Logit (Heuristic C) | 0.681 ± 0.002 | 0.561 ± 0.003 | 0.281 ± 0.01 | 0.539 ± 0.003 |
| | Stacked | 0.675 ± 0.004 | 0.548 ± 0.006 | 0.205 ± 0.023 | 0.535 ± 0.007 |

Table S4: Results of the in-sample evaluation (CV) of the 7 models for each SH-related characteristic. Values represent the mean and standard deviation across CV repetitions.

| SH Characteristic | Model | Test | p-value | p-value CI |
|---|---|---|---|---|
| **Daytime sleepiness** | Logit (Heuristic C) | partial | 0.0 | [0, 0.003] |
| | Logit (Heuristic C) | full | 0.0 | [0, 0.003] |
| | SVM (RBF Kernel) | partial | 0.0 | [0, 0.003] |
| | SVM (RBF Kernel) | full | 0.0 | [0, 0.003] |
| **Morning/Evening chronotype** | Linear SVM | partial | 0.754 | [0.726, 0.780] |
| | Linear SVM | full | 0.0 | [0, 0.003] |
| | Logit (Heuristic C) | partial | 0.153 | [0.131, 0.176] |
| | Logit (Heuristic C) | full | 0.0 | [0, 0.003] |
| | SVM (RBF Kernel) | partial | 0.19 | [0.166, 0.215] |

| | | | | |
|---|---|---|---|---|
| | SVM (RBF Kernel) | full | 0.0 | [0, 0.003] |
| **Easiness getting up in the morning** | Linear SVM | partial | 0.0 | [0, 0.003] |
| | Linear SVM | full | 0.0 | [0, 0.003] |
| **Daytime nap** | Logit (Heuristic C) | partial | 0.0 | [0, 0.003] |
| | Logit (Heuristic C) | full | 0.0 | [0, 0.003] |
| | SVM (RBF Kernel) | partial | 0.0 | [0, 0.003] |
| | SVM (RBF Kernel) | full | 0.001 | [2.5e-5, 0.005] |
| **Insomnia** | Linear SVM | partial | 0.0 | [0, 0.003] |
| | Linear SVM | full | 0.0 | [0, 0.003] |
| | Logit (Heuristic C) | partial | 0.0 | [0, 0.003] |
| | Logit (Heuristic C) | full | 0.0 | [0, 0.003] |
| | Stacked | partial | 0.0 | [0, 0.003] |
| | Stacked | full | 0.0 | [0, 0.003] |
| **Sleep duration** | Linear SVM | partial | 0.0 | [0, 0.003] |
| | Linear SVM | full | 0.0 | [0, 0.003] |
| | Logit (Heuristic C) | partial | 0.0 | [0, 0.003] |
| | Logit (Heuristic C) | full | 0.0 | [0, 0.003] |
| | Extra Trees | partial | 0.0 | [0, 0.003] |
| | Extra Trees | full | 0.0 | [0, 0.003] |
| **Snoring** | Linear SVM | partial | 0.0 | [0, 0.003] |
| | Linear SVM | full | 0.0 | [0, 0.003] |
| | Logit (Heuristic C) | partial | 0.0 | [0, 0.003] |
| | Logit (Heuristic C) | full | 0.0 | [0, 0.003] |

Table S5: Results of the confounding bias partial and full tests for the models that were selected as best performing at least for one of the 4 evaluated metrics.

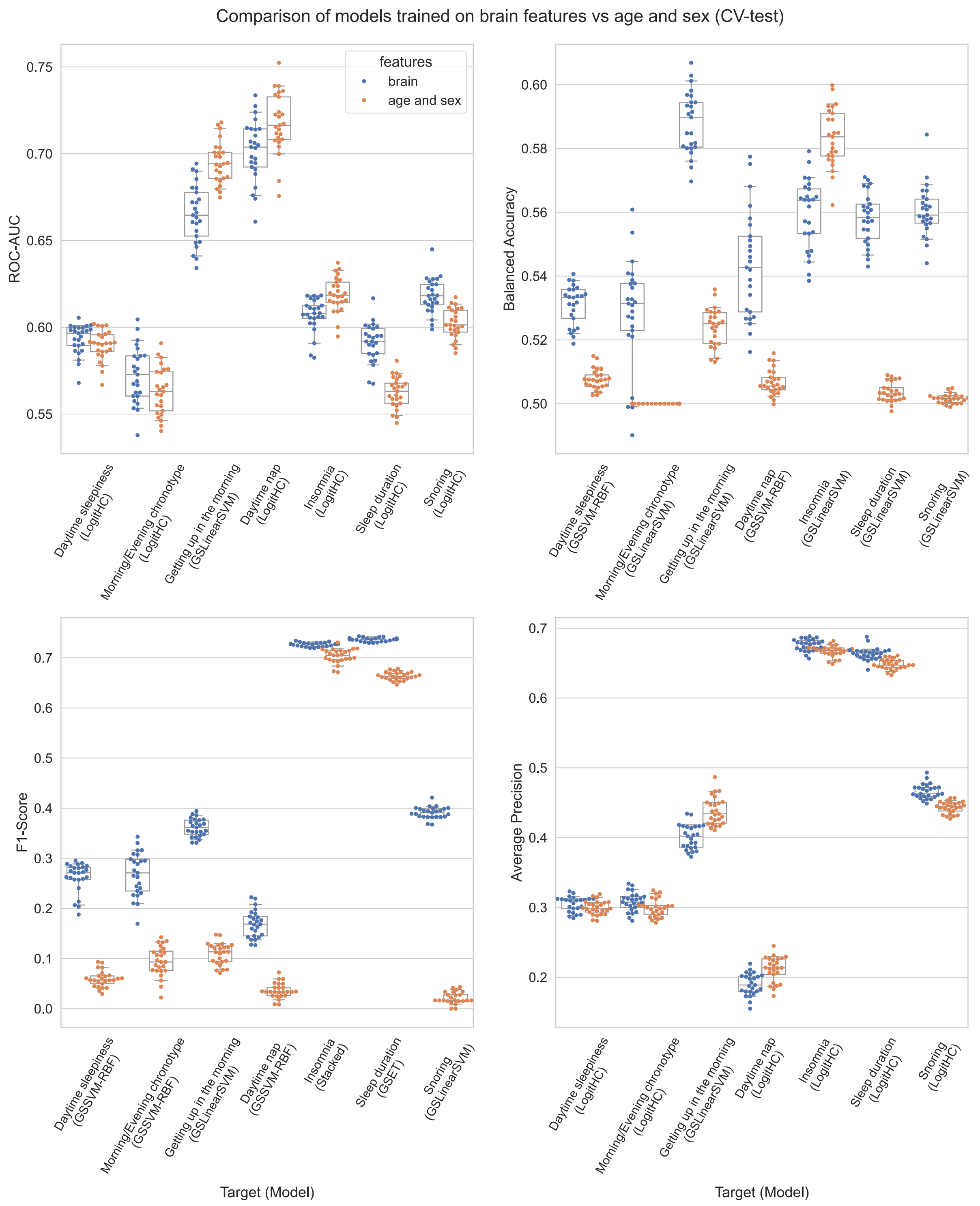


Figure S1: Side-by-side comparison of the best model for each SH-related characteristic using either brain features (blue) or age and sex (orange), for both threshold-dependent and threshold-independent metrics. Each dot represents the performance obtained at each of the 25 test folds within cross-validation (CV). Boxplots summarize the medians and 95% CI for the underlying distribution.

### References

1. R: Fast Heuristics For The Estimation Of the C Constant Of A... https://search.r-project.org/CRAN/refmans/LiblineaR/html/heuristicC.html.
